# Predicting the HER2 status in esophageal cancer from tissue microarrays using convolutional neural networks

**DOI:** 10.1101/2022.05.13.491769

**Authors:** Juan I. Pisula, Rabi R. Datta, Leandra Börner Valdez, Jan-Robert Avemarg, Jin-On Jung, Patrick Plum, Heike Löser, Philipp Lohneis, Monique Meuschke, Daniel Pinto dos Santos, Florian Gebauer, Alexander Quaas, Christiane J. Bruns, Axel Walch, Kai Lawonn, Felix C. Popp, Katarzyna Bozek

## Abstract

**Background:** Fast and accurate diagnostics are key for personalized medicine. Particularly in cancer, precise diagnosis is a prerequisite for targeted therapies which can prolong lives. In this work we focus on the automatic identification of gastroesophageal adenocarcinoma (GEA) patients that qualify for a personalized therapy targeting epidermal growth factor receptor 2 (HER2). We present a deep learning method for scoring microscopy images of GEA for the presence of HER2 overexpression.

**Methods:** Our method is based on convolutional neural networks (CNNs) trained on a rich dataset of 1,602 patient samples and tested on an independent set of 307 patient samples. We incorporated an attention mechanism in the CNN architecture to identify the tissue regions in these patient cases which the network has detected as important for the prediction outcome. Our solution allows for direct automated detection of HER2 in immunohistochemistry-stained tissue slides without the need for manual assessment and additional costly in situ hybridization (ISH) tests.

**Results:** We show accuracy of 0.94, precision of 0.97, and recall of 0.95. Importantly, our approach offers accurate predictions in cases that pathologists cannot resolve, requiring additional ISH testing. We confirmed our findings in an independent dataset collected in a different clinical center.

**Conclusions:** We demonstrate that our approach not only automates an important diagnostic process for GEA patients but also paves the way for the discovery of new morphological features that were previously unknown for GEA pathology.

## Background

Gastroesophageal adenocarcinoma (GEA) is the seventh most common cancer worldwide, with an increasing number of cases in the western hemisphere. Despite multimodal therapies with neoadjuvant chemotherapy/chemoradiation before surgery, median overall survival does not exceed four years [1–5]. Epidermal growth factor receptor 2 (HER2) encodes a transmembrane tyrosine kinase receptor and is present in different tissues e.g. epithelial cells, mammary gland, and the nervous system. It is also an important cancer biomarker. HER2 activation is associated with angiogenesis and tumorigenesis. Various solid tumors display HER2 overexpression, and targeted HER2 therapy improves their treatment outcomes [6]. Clinical guidelines for GEA recommend adding Trastuzumab – a monoclonal antibody binding to HER2 – to the first-line palliative chemotherapy for HER2-positive cases. HER2 targeting drugs are also currently investigated in the curative therapy for GEA [7].

Accurate testing for the HER2 status is a mandatory prerequisite for the application of targeted therapies. The gold standard for determining the HER2 status is an analysis of the immunohistochemical (IHC) HER2 staining by an experienced pathologist, if necessary followed by an additional in situ hybridization (ISH). The pathologist examines the immunohistochemistry staining of cancer tissue slides for HER2 and determines the IHC score ranging from 0 to 3. According to expert guidelines [8], the factors determining the score include the staining intensity, the number of connected positive cells, and the cellular location of the staining (Supplemental Table 1). The IHC scores 0 and 1 define patients with a negative HER2 status that are not eligible for targeted anti-HER2 therapy. An IHC score of 3 designates a positive HER2 status, and these patients receive Trastuzumab. A score of 2 is equivocal. In this case, an additional in situ hybridization (ISH) assay resolves the IHC score 2 as a positive or negative HER2 status. However, both manual scoring and additional ISH testing are time-consuming and costly.

Here, we ask whether Convolutional Neural Networks (CNNs) can directly predict the HER2 status from IHC-stained tissue sections without additional ISH testing. We explore a large tissue microarray (TMA) with 1,602 digitized images stained for HER2. We use this image dataset as a training set to train two different CNN classification models. We test these models on an independent test dataset of 307 TMA images from an unrelated patient group from the same center. We also further validate the HER2 status prediction accuracy of our approach on a patient cohort from a different clinical center. If successful, CNNs could assist pathologists in evaluating IHC stainings and, therefore, save time and expenses related to the ISH analysis.

## Methods

### Tumor sample and image preparation

For training the CNNs, we used a multi-spot tissue microarray (TMA) with 165 tumor cases and a single-spot TMA with 428 tumor cases, described elsewhere [9]. We additionally prepared an independent single-spot TMA with 307 tumor cases as the test dataset. The test set consisted of tumor cases that occurred at a later time point compared to the training set cases. This dataset construction strategy mimics how such a model would be developed and deployed in a clinical routine. Coincidentally, our test set does not include tumor cases with an IHC score of 1. The multi-spot TMA was composed of eight tissue cores (1.2 mm diameter) of each tumor - four cores punched on the tumor margin and four in the tumor center. To construct the single-spot TMA, we punched one tissue core per patient from the tumor center. The cores were transferred to TMA receiver blocks. Each TMA block contained 72 tissue cores. Subsequently, we prepared 4 μm-thick sections from the TMA blocks and transferred them to an adhesive-coated slide system (Instrumedics Inc., Hackensack, NJ).

We used a HER2 antibody (Ventana clone 4B5, Roche Diagnostics, Rotkreuz, Switzerland) on the automated Ventana/Roche slide stainer to perform immunohistochemistry (IHC) on the TMA slides. HER2 expression in carcinoma cells was assessed according to staining criteria listed in Table 1. Scores 0 and 1 indicated negative HER2 status, and score 3 indicated positive HER2 status. Immunohistochemical expression evaluation was assessed manually by two pathologists (A.Q. and H.L.) according to [10]. Discrepant results, which occurred only in a small number of samples, were resolved by consensus review. Spots with a score of 2 were analyzed by fluorescence in situ hybridization (FISH) to resolve the HER2 status. The FISH analysis evaluated the HER2 gene amplification status using the Zytolight SPEC ERBB2/CEN 17 Dual Probe Kit (Zytomed Systems GmbH, Germany) according to the manufacturer’s protocol. A fluorescence microscope (DM5500, Leica, Wetzlar, Germany) with a 63× objective was used for scanning the tumor tissue for amplification hotspots. We counted the signals in randomly chosen areas of homogeneously distributed signals. Twenty tumor cells were evaluated by counting green HER2 and orange centromere-17 (CEN17) signals. The reading strategy followed the recommendations of HER2/CEN17 ratio ≥ 2.0 or HER2 signals ≥ 6.0 for HER2 positive and a HER2/CEN17 ratio < 2.0 for HER2 negative samples.

**Table 1.** Results of the Attention Based MIL method on the tasks of IHC score prediction and HER2 status prediction.

| Task | Balanced<br>Acc. | Precision | Recall | F1 Score |
| --- | --- | --- | --- | --- |
| IHC score<br>prediction | 0.8249 | 0.9470 | 0.9185 | 0.9302 |
| HER2 status<br>prediction | 0.9429 | 0.9705 | 0.9478 | 0.9551 |

We digitized the slides with a digital slide scanner (NanoZoomer S360, Hamamatsu Photonics, Japan) with 40-times magnification and used QuPath’s [11] TMA dearrayer to slice the digitized slides into individual images (.jpg files, 5,468 × 5,468 pixels). After discarding corrupted images, this procedure yielded 1,602 images for training and validation, and 307 images for testing. Each image was labeled with the IHC score (0, 1, 2, or 3) and the HER2 status (0 or 1) that was determined by the pathologists or by FISH analysis in equivocal cases. This methodology corresponds to the gold standard, and we used this labeling in training the different machine learning models.

### Classification models

We used ResNet34 architecture [12] for prediction of IHC score and HER2 status. The network was trained as a four class IHC score classifier and as a binary classifier of the HER2 status. Given the large resolution of the tissue images (5,468 × 5,468 pixels), this approach required scaling them down by 5.34 to the size of 1,024 × 1,024 pixels to allow the network to train within our hardware memory limits.

The strongly reduced resolution of the downscaled microscopy images could potentially remove important visual details from the images that are key for the score assignment. To overcome this, we implemented a method that allows training neural networks on large images at their original resolution by exploiting weakly supervised Multiple Instance Learning (MIL) [13].

In the weakly supervised multiple instance learning approach, each slide is considered as a bag of smaller tiles (instances) whose respective individual labels are unknown. To make a bag-level prediction, image tiles are embedded in a low dimensional vector space, and the embeddings of individual tiles are aggregated to obtain representation of the entire image. This representation is used as input of a bag-level classifier.

For the aggregation of the tile embeddings, we used the attention-based operator proposed by Ilse et al. [14]. It consists of a simple feed-forward network that predicts an attention score for each of the embeddings. These scores indicate how relevant each tile is for the classification outcome, and are used to calculate a weighted sum of the tile representations as the aggregation operation. Weights of a bag sum to one, this way the bag representation is invariant to bag size. Finally, the bag vector representation is used as the input of a feed-forward neural network to perform the final classification.

In this approach, non-overlapping tiles of 224 × 224 pixels were extracted from each slide, and their embeddings were derived from a ResNet34 model. Empty tiles were discarded beforehand. As in the fully supervised approach, the MIL classifier was trained separately to predict IHC score and HER2 status.

We also constructed a method for predicting IHC score and HER2 status based on the staining intensity of the slides, a feature that is conventionally used by automatic IHC scoring software. This method was constructed to compare how predictive the single feature of staining intensity is compared to the higher level features learned by our CNN models. To extract the IHC staining expression from the images we used color deconvolution [15]. From the staining channel, non-overlapping tiles of 224 × 224 pixels were extracted and the average staining intensity was calculated for each tile. The staining intensity of each slide was then calculated as the maximum of the average intensities of its tiles. The proposed slide descriptor was used as input in two logistic regression classifiers to predict IHC score and HER2 status separately. This approach can also be seen as a multiple instance classification formulation where the feature extracted for each instance is its average staining intensity value, and the bag is aggregated using the maximum operator.

### Network training

The dataset showed an unbalanced distribution of the IHC score (Supplemental Fig. 1) reflecting the frequency of HER2 expression in the population [16]. To obtain representative training and validation sets, we split images of each IHC score in 80-20 proportions. For the samples with score 2, the 80-20 split was done separately for those with positive status and those with negative status. During training, we performed a weighted sampling of the images of each score such that each of the IHC scores is equally represented during training. We performed random horizontal and vertical flips as data augmentation.

We used Adam optimizer in training [17], with weight decay of 1×10^-8^ and betas of 0.9 and 0.999. The learning rates as well as their schedulers were chosen based on a hyperparameter search. The ResNet classifiers were trained using a learning rate 1×10^-5^, which was reduced by a factor of 0.1 if the accuracy of the validation set would not improve after 20 epochs of training. The MIL classifier was trained using a learning rate of 5×10^-9^, decreasing it by a factor of 0.3 if the accuracy of the validation set would not improve after 40 epochs. We used a batch size of 32 in the ResNet classifier and a batch size of only one full resolution image with a bag size depending on the amount of extracted tiles in the MIL classifier.

Computational work was performed on the CHEOPS high performance computer, on nodes equipped with 4 NVIDIA V100 Volta graphics processing units (GPUs). We used PyTorch (version 1.8.1) [18] for data loading, creating models, and training.

## Results

### IHC score prediction

First, we implemented a multiple instance learning (MIL) [13] method allowing us to make the classification of the images at their highest resolution. Using this technique the images are split into smaller tiles, encoded into their numeric embeddings and ranked using the attention mechanism as proposed by Ilse et al. [14]. The attention mechanism allows for automatic identification of areas in the image that are important for the predicted score, this way providing means to inspect and interpret the prediction outcomes of the network.

This technique has shown the balanced accuracy of 0.8249, precision of 0.9470 and recall of 0.9185 (Fig. 1:left, Table 1). Given the score imbalance and the lack of samples with an IHC score of 1 in the test set, the reported performance metrics were calculated in a balanced manner as an average of the metric of each individual label weighted by their number of samples of that given label. Most notably, the outermost classes 0 and 3 were predicted with the highest accuracy while ∼ 33% of score 2 images were incorrectly predicted.

**Figure 1.**
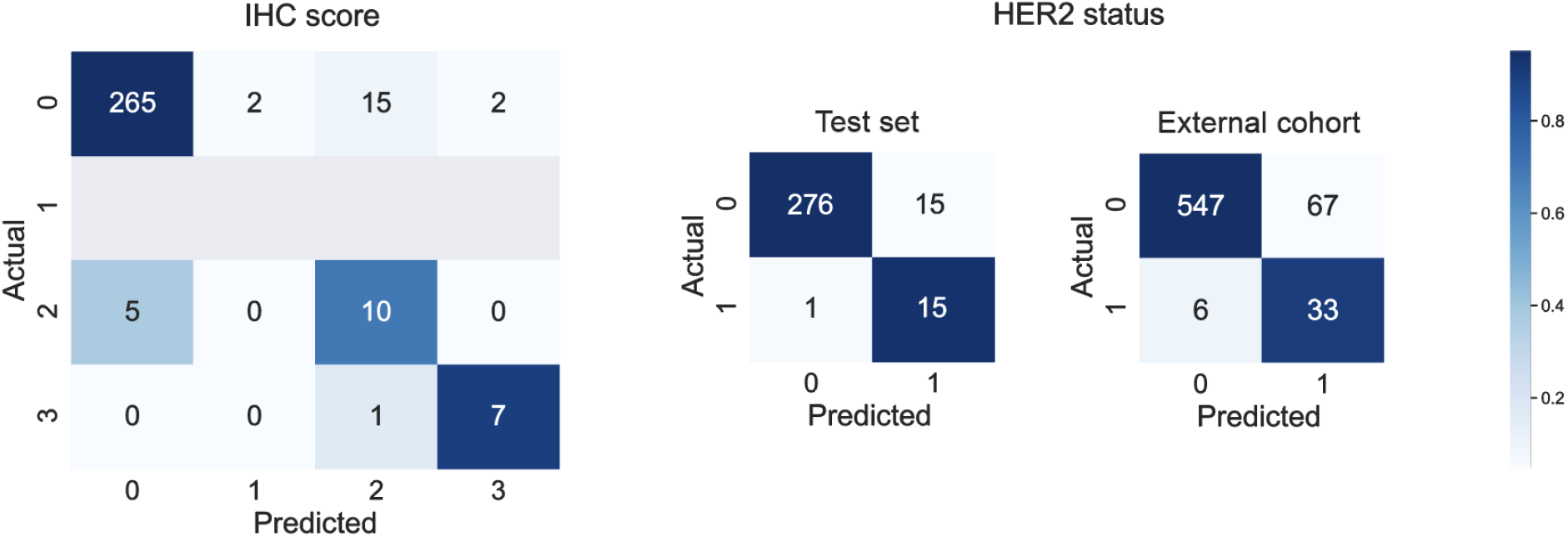
Confusion matrices of the IHC score prediction evaluated on the test set (left), and HER2 status prediction evaluated on the test set and on the external cohort (right).

We next examined whether a simpler CNN-based classification approach allows for predicting the IHC score from the TMA images. In order for these images to fit within our hardware constraints, we downsampled them by a factor of 5.34 to a size of 1,024 × 1,024 pixels. We trained classification architecture ResNet34 [12] on the rescaled dataset and analyzed it on the test set of images adjusted correspondingly. This approach resulted in balanced accuracy of 0.8536, precision of 0.9544 and recall of 0.8859 (Table 2). The almost equal accuracy and precision of this model suggests that relatively large visual details visible at a lower resolution are sufficient for the most accurate prediction.

**Table 2.** Cross-tabulation of true IHC score and predicted HER2 status of the test dataset. ‘2-’ and ‘2+’ scores stand for IHC score 2 and HER2 negative and positive status, respectively.

| Predicted HER2 status | True IHC score |  |  |  |
| --- | --- | --- | --- | --- |
|  | 0 | 2- | 2+ | 3 |
| Negative | 273 | 3 | 1 | 0 |
| Positive | 11 | 4 | 7 | 8 |

### HER2 status prediction

We next addressed the question whether the HER2 status can be predicted directly from the IHC-stained images, without additional ISH testing. Images in our dataset with IHC score of 0 or 1 are HER2 negative, those with a score of 3 are positive. Those with a score of 2 were additionally resolved using ISH resulting in the following positive/negative HER2 status split: 77/33% in the train set, 53/47% in the test set. Out of 15 IHC score 2 images in the test set, there were eight HER2 positive and seven HER2 negative. The train-validation split was done in such a way that all the score and status combinations are distributed equally in both sets.

The MIL classifier resulted in performance with balanced accuracy of 0.9429, precision of 0.9705 and recall of 0.94780 (Fig. 2, Table 2). As in the IHC score prediction task, the results were calculated as a weighted average of the individual metrics for class 0 (HER2 negative) and class 1 (HER2 positive) to take account of the class imbalance. Within both the HER2 negative and HER2 positive classes, less than 7% of images were misclassified resulting in balanced precision and recall > 0.94. To better understand the errors of the model, we additionally inspected the HER2 status prediction accuracy within images of different IHC scores (Table 3). With ∼27% false positive and ∼7% false negative predictions, the highest error rate occurred in images with the IHC score of 2. The higher proportion of false positives among the score 2 images could be due to the underrepresentation of samples with this IHC score and negative HER2 status in the training set in the score 2 images. In images with IHC scores 0 and 3, the prediction error was below 4%. The difference in performance between the 4-class and the binary classifiers suggests that the inter-score differences are more subtle than the ones differentiating the two HER2 statuses.

**Figure 2.**
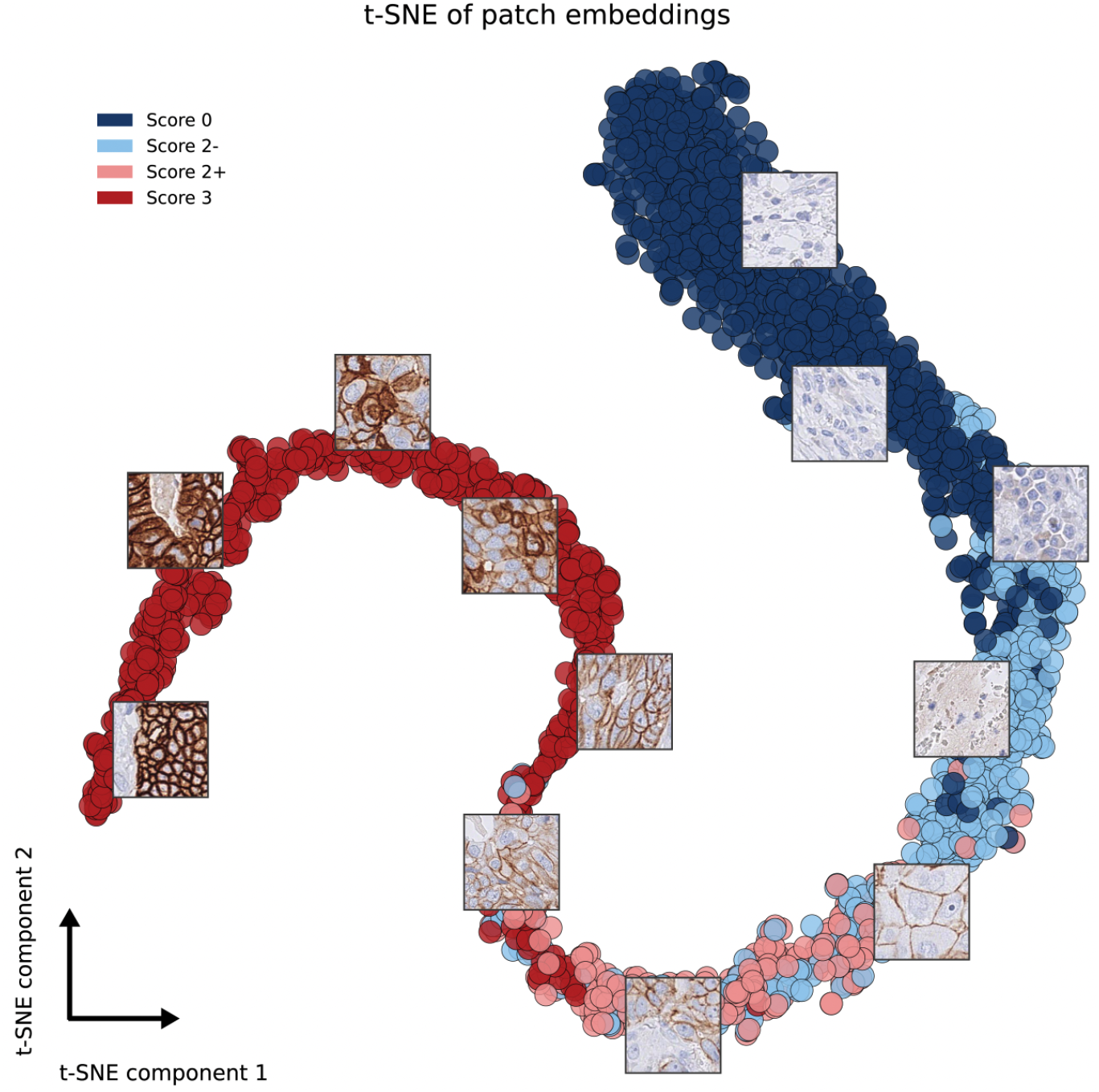
t-SNE visualization of tile embeddings produced by the IHC score MIL classifier on the test set images, with the vectors coloured according to the score of their respective slides. Visual similarity of the tiles is reflected in their neural network-derived representations and the embeddings of similar tiles are close in the learned vector space. Coincidentally, there are no TMA images with a score of 1 in the test set because the test set consisted of the consecutive tumor cases that followed the training set cases.

### Performance on external cohort

Even if independently, our train and test datasets were collected and prepared within one hospital. To verify how the performance of our model is dependent on the aspects related to the data preparation, we evaluated our models on an independent cohort from a different clinical center [20]. In particular we aimed to investigate whether HER2 status prediction is indeed possible using IHC-stained images only. The external cohort included 653 tissue samples belonging to 297 patients with the following IHC score distribution: 416/186/14/37 samples of scores 0/1/2/3 respectively. Out of the score 2 samples, 12 showed a negative HER2 status and 2 samples showed positive HER2 status.

Given the different color distribution and potential staining quality deterioration due to the sample age, we applied a preprocessing step to these images. We used Macenko’s method for stain estimation [21] together with color deconvolution/convolution [15] to match the staining to our in-house dataset.

The MIL classifier yielded a balanced accuracy of 0.8688, precision of 0.9490 and recall of 0.8908 (Fig. 1). These results support the applicability of our approach in an important clinical context where the distinction of HER2 status is key for further treatment.

### Insights into the learning process of the MIL classifier

The ResNet and the MIL classifiers achieved almost identical accuracy on our in-house test set in both the IHC score and the HER2 status prediction. However, the advantage of the more compute-intensive weakly supervised MIL approach is the possibility to inspect the visual features that the network utilizes in the classification process. The embeddings and attention scores assigned to individual 224 × 224 pixel tiles can provide insights into the key visual features used by the MIL approach in the classification.

First, we examined via t-distributed stochastic neighbor embedding (tSNE) dimensionality reduction method [19] the embeddings of the image tiles in the test set generated by the IHC score prediction network (Fig. 2). In this visualization, the spatial proximity of tiles reflects the similarity of their embeddings. Although the network was trained on the IHC score, it also correctly separates the HER2 status of the parent TMA image. HER2 negative tiles with a score of 2 (2-) group together with score 0 tiles, and HER2 positive tiles with a score of 2 (2+) group together with score 3 tiles.

Additionally, neighboring tiles in the tSNE projection show visual similarity. Most strikingly, tiles grouped together show a similar staining intensity and this intensity gradually changes along the 2D projection of the embeddings. Staining intensity is, however, not the only visual feature determinant of the HER2 scoring, which also takes additional morphological features into account (Table 1). We expect these morphological features to also be encoded in the learned vector space.

Next, we inspected the attention score of the MIL classifier and its distribution within the tissue slides. The attention value reflects the importance of a given image tile for the final prediction score and this way provides information on spatial distribution of the visual features in the tissue that the network is exploiting in the prediction. Since the IHC staining is insufficient to resolve the HER2 status if the tissue IHC score is 2, we inspected which visual features are exploited by the network in resolving the HER2 status of the score 2 tissue slides (Fig. 3). Strikingly, the attention of the MIL classifier for the HER2 status focuses on areas of high staining intensity and corresponds to the mean intensity of the tiles at first sight.

**Figure 3.**
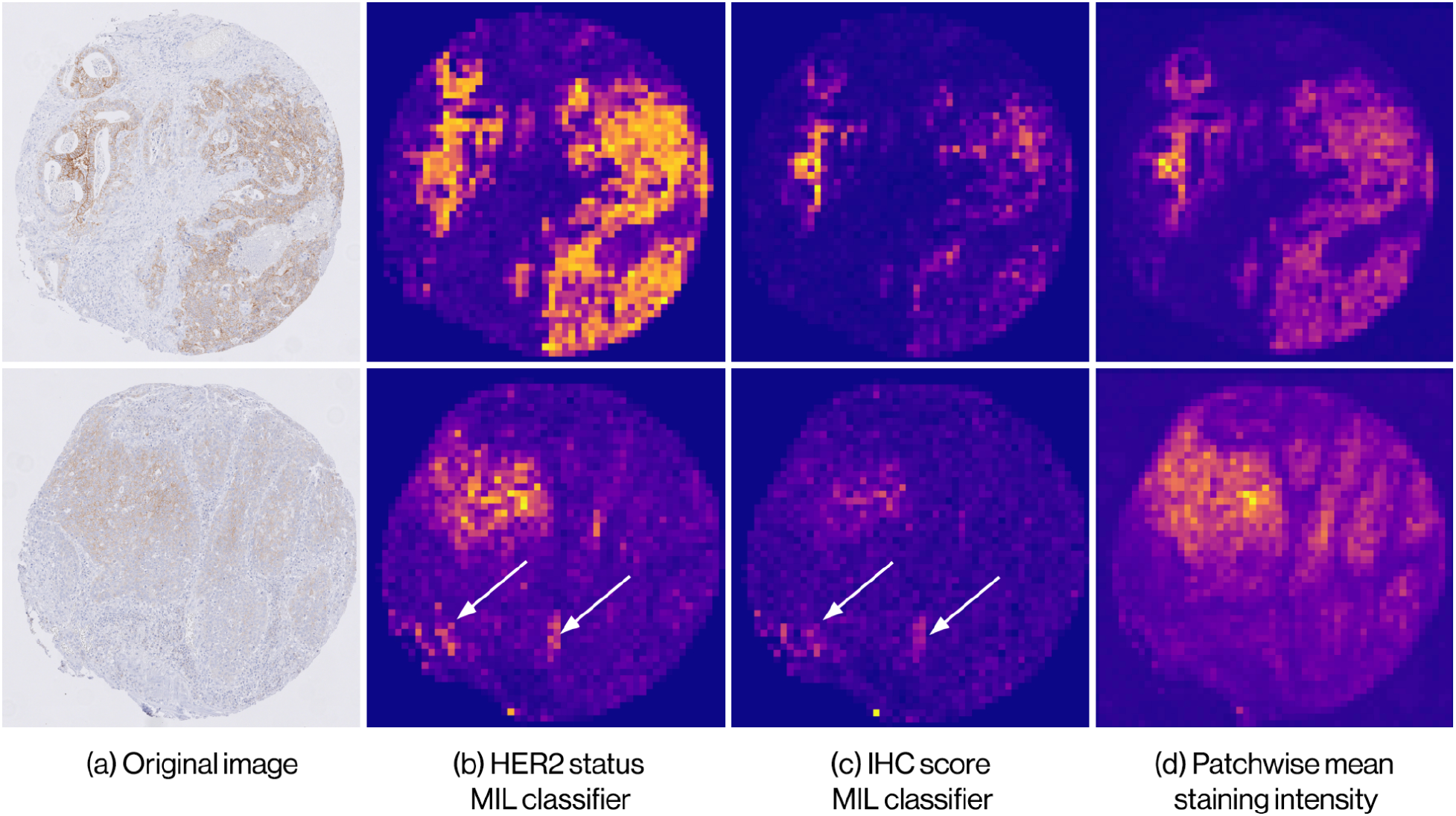
Heatmap visualizations of the attention value and mean staining intensity in tiles within the tissue image. a) Slides with IHC score 2 and negative HER2 status. b) Attention score heatmap of HER2 status MIL classifier. c) Attention score heatmap of IHC score MIL classifier. d) Patchwise mean staining intensity heatmap. White arrows point to locations where the attention values do not match staining intensity.

Given the relationship of the embeddings as well as attention value to the staining intensity, we tested the accuracy of a predictive model based on the staining intensity only. Similar to the tiling approach of the MIL classifier, we split the tissue slides in 224 × 224 pixel tiles and averaged the staining intensity in each of the tiles. We, then, used the maximum of the average intensities across the tiles of an image as the quantitative descriptor of the entire image. We trained two logistic regression models to predict IHC score and HER2 status, respectively. The stain intensity-based model showed a balanced accuracy of 0.6876 in the prediction of the IHC score, markedly lower compared to the MIL classifier with a balanced accuracy of 0.8249. The major difference in performance between these models is in images with an IHC score of 2 (Fig. 4). In the task of predicting the HER2 status, the balanced accuracy of the staining intensity-based model reached 0.8457 compared to 0.9429 of the MIL classifier. These results suggest that not only the staining intensity but also additional morphological features are considered by the deep learning models in the classification. These features are particularly important for correct recognition of images belonging to the intermediate IHC score 2. We indicate examples of such features in Figure 3. Even though attention value and staining intensity largely match, the heatmaps in Figure 3 demonstrate prominent exceptions where features of high attention do not show high staining intensity.

**Figure 4.**
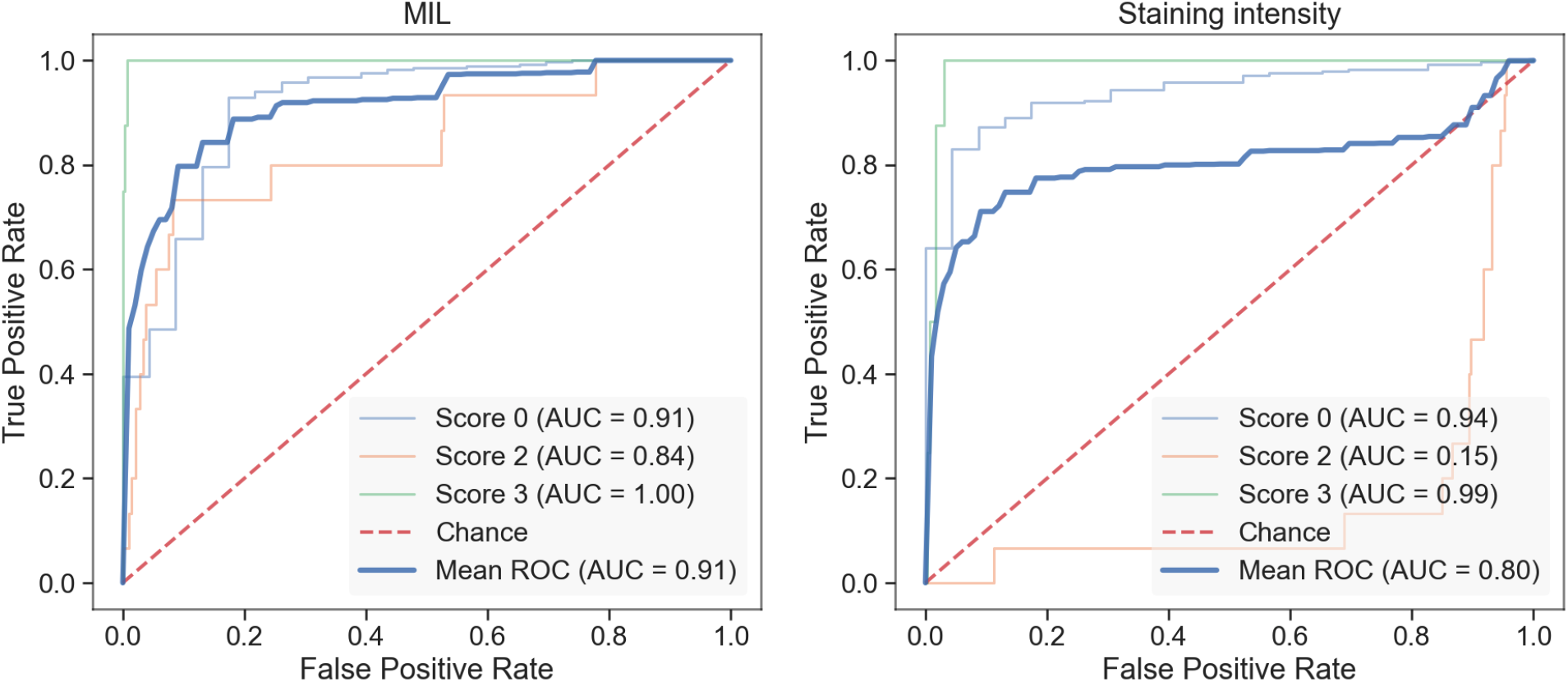
Per-class ROC curves for the IHC score classifiers, calculated in a “one-vs-all” fashion of the MIL (left panel) and staining intensity-based classifier (right panel). While both models’ performance is similar for images of score 0 and 3, images of score 2 are not possible to correctly recognize based on staining intensity only.

## Discussion

Automated and accurate image-based diagnostics help to accelerate medical treatment and decrease the work burden of the medical personnel. Here, we demonstrate that deep learning-based prediction of the IHC score (0 - 3) and the HER2 status (negative or positive) is generally possible with a balanced accuracy of ∼0.85 and ∼0.94, respectively. Among the scores, IHC score 2 images show the highest proportion of misclassified samples. These score 2 images cannot be unequivocally classified regarding their HER2 status by the pathologists and need further ISH staining. While it is considered that it is not possible to resolve the HER2 status based on the IHC staining of the IHC score 2 images, our models correctly predict the HER2 status of 73% of these images in our test dataset. Notably, score 2 samples are strongly underrepresented in our datasets. We expect that with more training samples of the underrepresented scores this prediction accuracy will improve.

We demonstrate that not only staining intensity - conventionally used in automated prediction tools - but also additional morphological properties are taken into account by the neural networks in the classification. The attention maps of the MIL classifier do not match the staining intensity. Additionally, the prediction based on the intensity only yields markedly lower accuracy. Unexpectedly however, the classifiers based on low- (1,024 × 1,024 pixel) and high-(5,468 × 5,468 pixel) resolution images achieve matched accuracy. Potentially, the lower resolution used in this study is sufficient to encode the key morphological features of the images. This resolution was the highest that still allowed for training ResNet within our hardware memory. Notably decreasing the size of the images further to 512 × 512 pixel size resulted in the decrease of the model balanced accuracy to 0.8200 for the prediction of IHC score. Unlike in this study, whole slide images (WSIs) instead of TMAs are used in the diagnostic pathological assessment. The WSI size is several orders of magnitude larger than the images in our dataset which does not allow for using simple classification architectures such as ResNet and MIL approaches are typically used instead. Our results suggest that reducing image resolution even 5-fold does not affect the deep learning model performance which could accelerate model training and reduce computational costs of models built on WSIs without compromising their accuracy.

Given the class imbalance of our datasets, we report the balanced accuracy and weighted recall, precision and F1 metrics as described previously, as the unbalanced and unweighted metrics may be misleading in describing the performance of the models. As an example, if unbalanced, the accuracy score of an IHC score classifier that always predicts score 0 would be 0.92 in our dataset, and an analogous HER2 status classifier would achieve accuracy of 0.94. Similarly, the unbalanced precision (and subsequently, F1) of our HER2 status classifiers would be inaccurate. If we take, for example, the MIL HER2 status classifier, its unbalanced precision score is 0.51, while its false positive rate is only 0.04. For these reasons we calculate our accuracy metrics in a balanced manner.

We propose that artificial intelligence-based HER2 status evaluation represents a valuable tool to assist clinicians. In particular, the attention map generated by the MIL classifier can aid the pathologists in their daily work by indicating the image areas of high information content for the evaluation. This approach could facilitate and speed up the manual analysis of large tissue images. The IHC score determination network can easily be transferred to any IHC staining other than HER2, further paving the way for digital pathology. We additionally demonstrate the capacity of our method to perform on samples from external clinical centers with similar prediction accuracy. We expect the power and generalizability of our deep learning model to increase with larger, multi-center datasets.

Finally, the high performance of our models in predicting the HER2 status of score 2 samples for which the status is considered as unresolvable based on the IHC staining, suggests that there exist visual features to be predictive of the HER2 status in these images as well. While identification of these features would require more IHC score 2 image data than available in our dataset, we expect that further deployment of the MIL models might lead to the discovery of novel morphological features improving image-based diagnostics.

## Conclusion

We demonstrate that it is possible to automatically predict HER2 overexpression directly from immunohistochemistry-stained images of gastroesophageal cancer tissue, an important diagnostic process in the treatment of GEA patients. CNNs not only replicate the IHC scoring system used by pathologists, but can directly predict HER2 status in cases where it is considered not possible to resolve this condition by IHC staining alone.

Staining intensity is not the only predictive feature of HER2 overexpression in the IHC images, and deep learning algorithms are able to capture more complex visual information such as morphological features present in these images. Furthermore, we find that the performance of models that use downscaled versions of our images achieve a similar performance than that of the models that make use of the full resolution data, meaning that such fine visual detail is not required to capture the important semantic information of the images.

We conclude that deep learning based image analysis represents a valuable tool both for the development of useful digital pathology applications and the discovery of visual features and patterns previously unknown to traditional pathology.

## Acknowledgments

Both K.B. and J.I.P. were hosted by the Center for Molecular Medicine Cologne throughout this research. K.B. and J.I.P. were supported by the BMBF program Junior Group Consortia in Systems Medicine (01ZX1917B) and BMBF program for Female Junior Researchers in Artificial Intelligence (01IS20054). This study was uploaded to bioRxiv as a preprint.

## Authors’ contribution

J.I.P. performed data analysis, paper writing and software development. R.R.D., L.B.V., J.J., P.P., P.H., M.M., D.P.S. and K.L. did the data acquisition. J.R.A. contributed to software development. H.K., F.G., A.Q. performed data acquisition and analysis. C.J.B. and K.L. revised the paper. A.W. provided the external cohort. F.C.P. and K.B. designed the study and performed data analysis and writing of the paper.

## Ethics approval and consent to participate

The study was performed in accordance with the ethical standards of the institutional research committee and with the 1964 Helsinki declaration and its later amendments. The present study was approved by the ethics committee of the University of Cologne (reference no. 13-091). Written informed consent was obtained from all patients.

## Consent for publication

Not applicable.

## Data availability

The data that support the findings of this study are available upon request from the corresponding author.

## References

1. Dai T, Shah MA. Chemoradiation in oesophageal cancer. Best Practice & Research Clinical Gastroenterology. 2015 Feb 1;29(1):193–209.

2. van Hagen P, Hulshof MC, Van Lanschot JJ, Steyerberg EW, Henegouwen MV, Wijnhoven BP, Richel DJ, Nieuwenhuijzen GA, Hospers GA, Bonenkamp JJ, Cuesta MA. Preoperative chemoradiotherapy for esophageal or junctional cancer. New England Journal of Medicine. 2012 May 31;366(22):2074–84.

3. Xi M, Hallemeier CL, Merrell KW, Liao Z, Murphy MA, Ho L, Hofstetter WL, Mehran R, Lee JH, Bhutani MS, Weston B. Recurrence risk stratification after preoperative chemoradiation of esophageal adenocarcinoma. Annals of surgery. 2018 Aug 1;268(2):289–95.

4. Noordman BJ, Verdam MG, Lagarde SM, Hulshof MC, Hagen PV, van Berge Henegouwen MI, Wijnhoven BP, van Laarhoven HW, Nieuwenhuijzen GA, Hospers GA, Bonenkamp JJ. Effect of neoadjuvant chemoradiotherapy on health-related quality of life in esophageal or junctional cancer: results from the randomized CROSS trial. Journal of Clinical Oncology. 2018 Jan 20;36(3):268–75.

5. Shapiro J, Van Lanschot JJ, Hulshof MC, van Hagen P, van Berge Henegouwen MI, Wijnhoven BP, van Laarhoven HW, Nieuwenhuijzen GA, Hospers GA, Bonenkamp JJ, Cuesta MA. Neoadjuvant chemoradiotherapy plus surgery versus surgery alone for oesophageal or junctional cancer (CROSS): long-term results of a randomised controlled trial. The lancet oncology. 2015 Sep 1;16(9):1090–8.

6. Oh DY, Bang YJ. HER2-targeted therapies—a role beyond breast cancer. Nature Reviews Clinical Oncology. 2020 Jan;17(1):33–48.

7. Wagner AD, Grabsch HI, Mauer M, Marreaud S, Caballero C, Thuss-Patience P, Mueller L, Elme A, Moehler MH, Martens U, Kang YK. EORTC-1203-GITCG-the “INNOVATION”-trial: Effect of chemotherapy alone versus chemotherapy plus trastuzumab, versus chemotherapy plus trastuzumab plus pertuzumab, in the perioperative treatment of HER2 positive, gastric and gastroesophageal junction adenocarcinoma on pathologic response rate: a randomized phase II-intergroup trial of the EORTC-Gastrointestinal Tract Cancer Group, Korean Cancer Study Group and Dutch Upper GI-Cancer group. BMC cancer. 2019 Dec;19(1):1–9.

8. Nie J, Lin B, Zhou M, Wu L, Zheng T. Role of ferroptosis in hepatocellular carcinoma. Journal of cancer research and clinical oncology. 2018 Dec;144(12):2329–37.

9. Plum PS, Gebauer F, Krämer M, Alakus H, Berlth F, Chon SH, Schiffmann L, Zander T, Büttner R, Hölscher AH, Bruns CJ. HER2/neu (ERBB2) expression and gene amplification correlates with better survival in esophageal adenocarcinoma. BMC cancer. 2019 Dec;19(1):1–9.

10. Lordick F, Al-Batran SE, Dietel M, Gaiser T, Hofheinz RD, Kirchner T, Kreipe HH, Lorenzen S, Möhler M, Quaas A, Röcken C. HER2 testing in gastric cancer: results of a German expert meeting. Journal of cancer research and clinical oncology. 2017 May;143(5):835–41.

11. Bankhead P, Loughrey MB, Fernández JA, Dombrowski Y, McArt DG, Dunne PD, McQuaid S, Gray RT, Murray LJ, Coleman HG, James JA. QuPath: Open source software for digital pathology image analysis. Scientific reports. 2017 Dec 4;7(1):1–7.

12. He K, Zhang X, Ren S, Sun J. Deep residual learning for image recognition. InProceedings of the IEEE conference on computer vision and pattern recognition 2016 (pp. 770–778).

13. Dietterich TG, Lathrop RH, Lozano-Pérez T. Solving the multiple instance problem with axis-parallel rectangles. Artificial intelligence. 1997 Jan 1;89(1-2):31–71.

14. Ilse M, Tomczak J, Welling M. Attention-based deep multiple instance learning. InInternational conference on machine learning 2018 Jul 3 (pp. 2127–2136). PMLR.

15. Ruifrok AC, Johnston DA. Quantification of histochemical staining by color deconvolution. Analytical and quantitative cytology and histology. 2001 Aug 1;23(4):291–9.

16. Koopman T, Smits MM, Louwen M, Hage M, Boot H, Imholz AL. HER2 positivity in gastric and esophageal adenocarcinoma: clinicopathological analysis and comparison. Journal of cancer research and clinical oncology. 2015 Aug;141(8):1343–51.

17. Kingma DP, Ba J. Adam: A method for stochastic optimization. arXiv preprint 1412.6980. 2014 Dec 22.

18. Paszke A, Gross S, Massa F, Lerer A, Bradbury J, Chanan G, Killeen T, Lin Z, Gimelshein N, Antiga L, Desmaison A. Pytorch: An imperative style, high-performance deep learning library. Advances in neural information processing systems. 2019;32.

19. Van der Maaten L, Hinton G. Visualizing data using t-SNE. Journal of machine learning research. 2008 Nov 1;9(11).

20. Langer R, Rauser S, Feith M, Nährig JM, Feuchtinger A, Friess H, Höfler H, Walch A. Assessment of ErbB2 (Her2) in oesophageal adenocarcinomas: summary of a revised immunohistochemical evaluation system, bright field double in situ hybridisation and fluorescence in situ hybridisation. Modern Pathology. 2011 Jul;24(7):908–16.

21. Macenko M, Niethammer M, Marron JS, Borland D, Woosley JT, Guan X, Schmitt C, Thomas NE. A method for normalizing histology slides for quantitative analysis. In2009 IEEE international symposium on biomedical imaging: from nano to macro 2009 Jun 28 (pp. 1107–1110). IEEE.

