## Supplemental material for "Predicting the HER2 status in esophageal cancer from tissue microarrays using convolutional neural networks"

**Supplemental Table 1**. Staining patterns used by pathologists to assess the IHC score of HER2 stainings in biopsies. This analysis method was used because TMAs resemble biopsies more than whole slides.

| IHC score | Pattern of IHC staining for HER2 | HER2 status |
|---|---|---|
| **0** | No reactivity or membranous reactivity in any (or <5) tumor cell(s) | ***negative*** |
| **1** | Tumor cell cluster with a very weak membranous reactivity (at least 5 tumor cells) | ***negative*** |
| **2** | Tumor cell cluster with a weak to moderate complete, basolateral or lateral only membranous reactivity (at least 5 tumor cells) | ***equivocal* (ISH assessment required)** |
| **3** | Tumor cell cluster with a strong complete, basolateral or lateral only membranous reactivity (at least 5 tumor cells) | ***positive*** |

**Supplemental Figure 1**. IHC score distribution of our in-house datasets (with score 2 separated by positive and negative HER2 status).

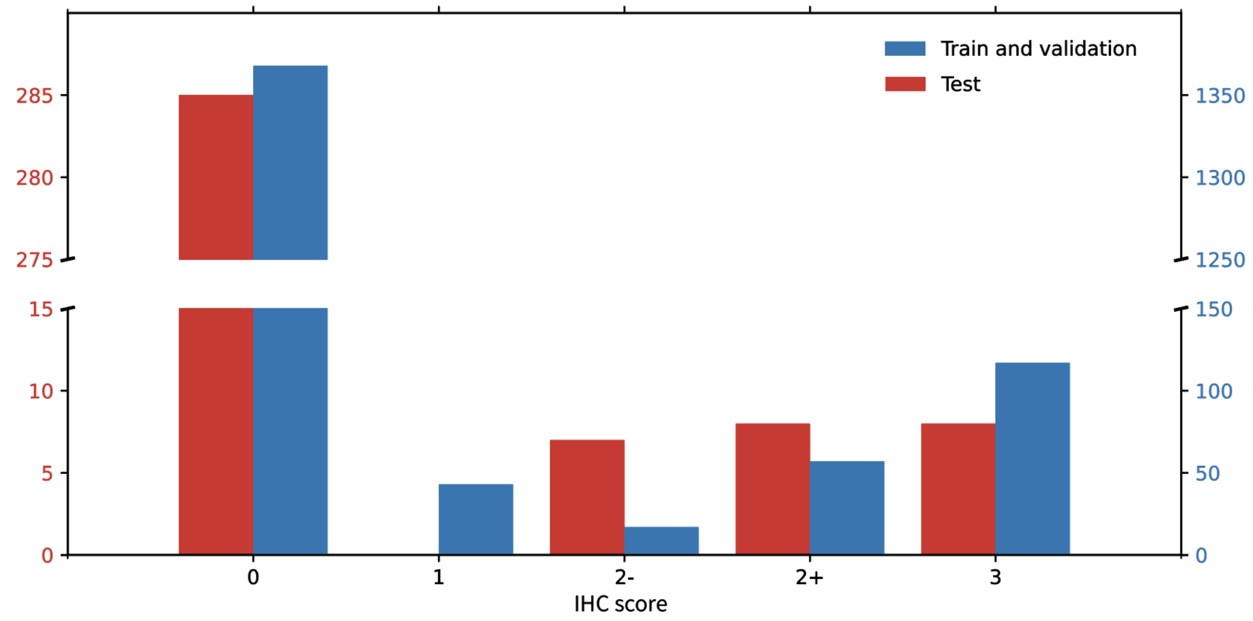
